# Drug repurposing high-throughput screen identifies candidate antiviral compounds against Puumala Orthohantavirus

**DOI:** 10.64898/2026.03.23.713563

**Authors:** Wanda Christ, Bartlomiej Porebski, Oscar Fernandez-Capetillo, Jonas Klingström

## Abstract

Hantaviruses are zoonotic negative-sense RNA viruses that cause two severe diseases; haemorrhagic fever with renal syndrome (HFRS) and hantavirus pulmonary syndrome (HPS) for which no approved antiviral therapies are available. To identify host-directed modulators of hantavirus infection in the available annotated drug space, we performed a drug repurposing screen in A549 cells and HUVECs, using live Puumala virus (PUUV). We identified and validated 70 drugs with antiviral activity across these 2 different cell systems. Functional clustering confirmed the known infection-inhibitory effect of several group of compounds, including inhibitors of heat shock proteins, mTOR pathway or nucleotide synthesis. In addition, we also identified compounds yet unexplored as antivirals against Hantaviruses, such as certain antibiotics. This dataset provides a systematic map of host pathways influencing PUUV infection and highlights candidate compounds and cellular processes that warrant further investigation.

## Introduction

Orthohantaviruses (hereafter referred to as hantaviruses) are zoonotic, negative-sense RNA viruses of the *Hantaviridae* family carried by rodents and other small animals[1]. Of the >60 hantaviruses identified to date several rodent-borne hantaviruses are associated with haemorrhagic fever with renal syndrome (HFRS) or hantavirus pulmonary syndrome (HPS)[2]. Annual incidence and case fatality vary by viral species and geographic region[3]. HFRS occurs predominantly in China, South Korea and parts of Europe, with case fatality rates normally below or around <1% but reaching up to 10% for certain hantaviruses[3]. Puumala virus (PUUV) is the most common hantavirus in Europe, where it causes HFRS with <1% case fatality. In contrast, HPS is endemic to South and North America and is associated with case fatality rates of 30 to 40%[3–7] .

To date, no EMA- or FDA-approved vaccine or hantavirus-specific treatment is available, and the treatment of hantavirus disease is limited to symptoms management[3,8] . Over the past decades, hypothesis-driven studies have proposed several antiviral strategies that showed promise in vitro or in animal models but have failed to demonstrate clear benefits in clinical settings (reviewed in[9,10] ). Reported drug discovery efforts have relied on pseudotyped viruses[11,12] or focused on specific facets of infection[13,14] . While informative, such approaches do not account for possible host dependencies in the context of productive live-virus infection and replication. To address these limitations, we investigated the Drug Repurposing Hub library[15] using a high-throughput screening strategy based on live virus, in search for possible novel antivirals. This approach enabled systematic identification of host-directed modulators of PUUV infection.

## Results

### A high-throughput screening platform for detection of PUUV-infected cells

To identify new antiviral compounds against hantavirus infection, we established a phenotypic screening assay based on PUUV infection of A549 cells and high-throughput immunofluorescence microscopy (**Figure 1A-B**). To maximise the performance and enable automation, we simplified the infection protocol, optimised the virus dose, and reduced sample preparation steps for microscopy without compromising staining quality (**Figure S1A-C**). The screening assay showed a reproducible infection rate of about 20% with low variation, providing a good screening window (Z’ = 0.7, S/N = 42). The assay was sensitive enough to detect both mild and strong antiviral effects of 2 tested control compounds, Ribavirin[16] and Bafilomycin A1[17] (**Figure 1C**).

**Figure 1.**
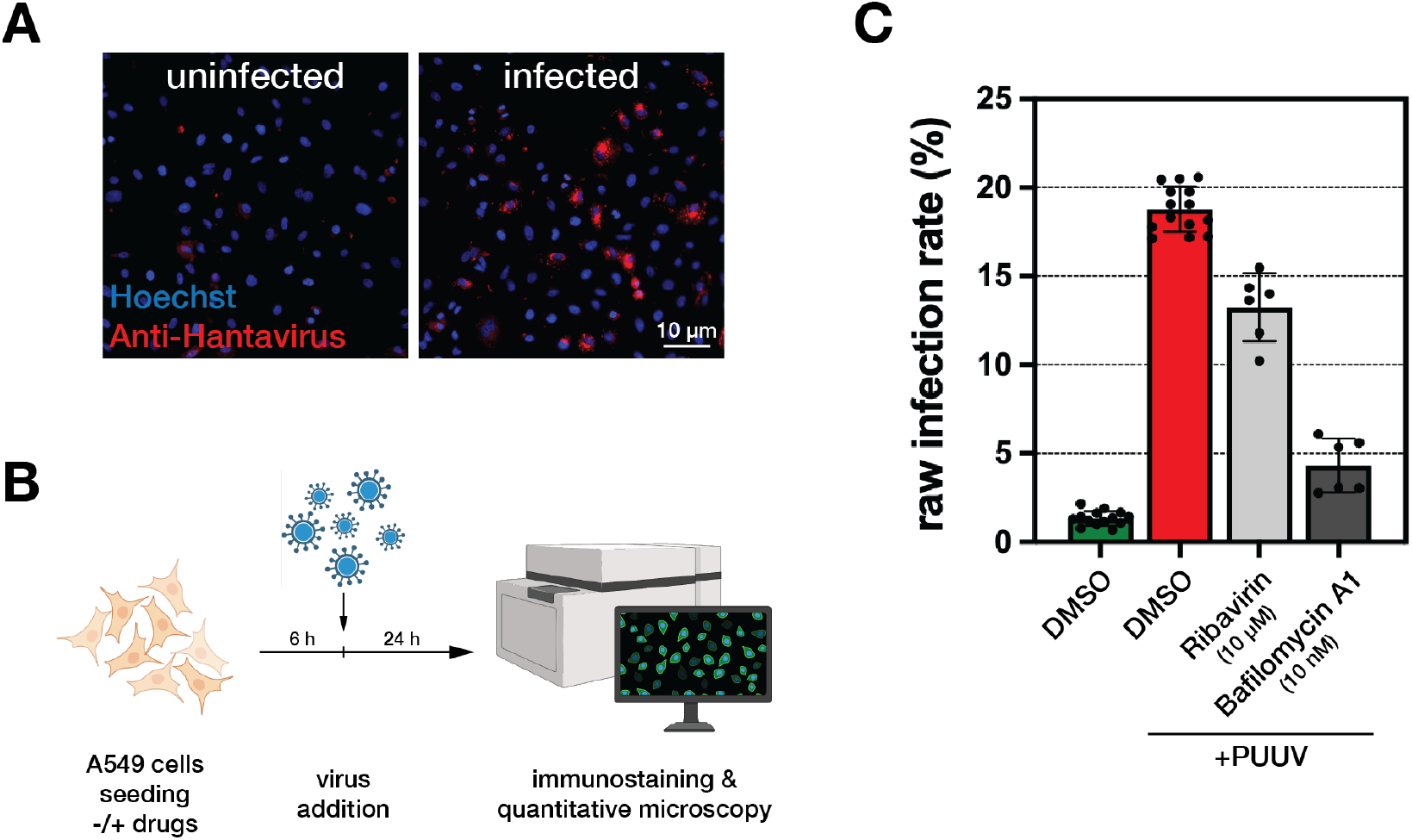
Screening assay development. A. Representative microscopic images of uninfected (left) or PUUV-infected (right) A549 cells. Viral antigen was detected using polyclonal antibody from convalescent sera (red), and nuclei were counterstained with Hoechst 3342 (blue). Scale bar, 10 µm. B. Schematic of the screening assay. A549 cells were seeded together with small molecule compounds and incubated for 6 hours. PUUV was then added, and 24 hours post infection cells were fixed and subjected to immunostaining and quantitative microscopy. C. Quantification of PUUV infection in A549 cells following treatment with vehicle (DMSO), 10 µM Ribavirin, or 10 nM Bafilomycin A1. Infection rate was calculated as the percentage of virus-positive cells. Dots represent individual wells, each with 2000-2500 cells. Bars indicate mean ± SD.

### Drug repurposing chemical screen for compounds with antiviral properties

Having established the screening assay, we next performed a screen, using the Drug Repurposing Hub library, a curated collection of 5256 FDA-approved, clinical, and pre-clinical drugs, to identify compounds that affect PUUV-infection. In the screening pipeline, A549 cells were seeded onto 384-well plates with pre-spotted library compounds (10 µM, triplicate plates), followed by the addition of PUUV. Twenty-four hours after infection, cells were fixed, immunostained with serum against viral proteins, and counter-stained with Hoechst 3342 and CellTracker to mark DNA and cell body, respectively. Samples were then imaged with high-throughput high-content automated microscopy and images were analysed in Cell Profiler[18] . Infection rates were calculated as the ratio of virus signal-positive cells to total number of cells. All data were normalised to the infection rate observed in vehicle (DMSO)-treated samples. In parallel, viability was calculated as the vehicle-normalised nuclei count. Data normalisation was done within each plate to account for plate-to-plate and batch-to-batch variation. Across the full screen, the mean infection rate was 17.7% with limited batch-to-batch variation (**Figure 2A and Figure S2A**). The Z’ across the full screen was 0.41 (**Figure S2B**), reflecting increased biological variability at scale but remaining within an acceptable range for phenotypic screening. For hit calling, we selected two criteria: compounds had to decrease the infection rate at least two-fold and show maximum 50% viability decrease (**Figure 2B**). One hundred fifty compounds passed these thresholds (**Figure 2C**). In addition, a substantial number of compounds led to an increase in the number of infected cells, with 25 passing the applied thresholds of viability and 2-fold increase in the infection rate (**Figure 2D**). Together, the screen demonstrated stable performance at scale and identified 150 antiviral and 25 proviral candidate compounds for subsequent validation.

**Figure 2.**
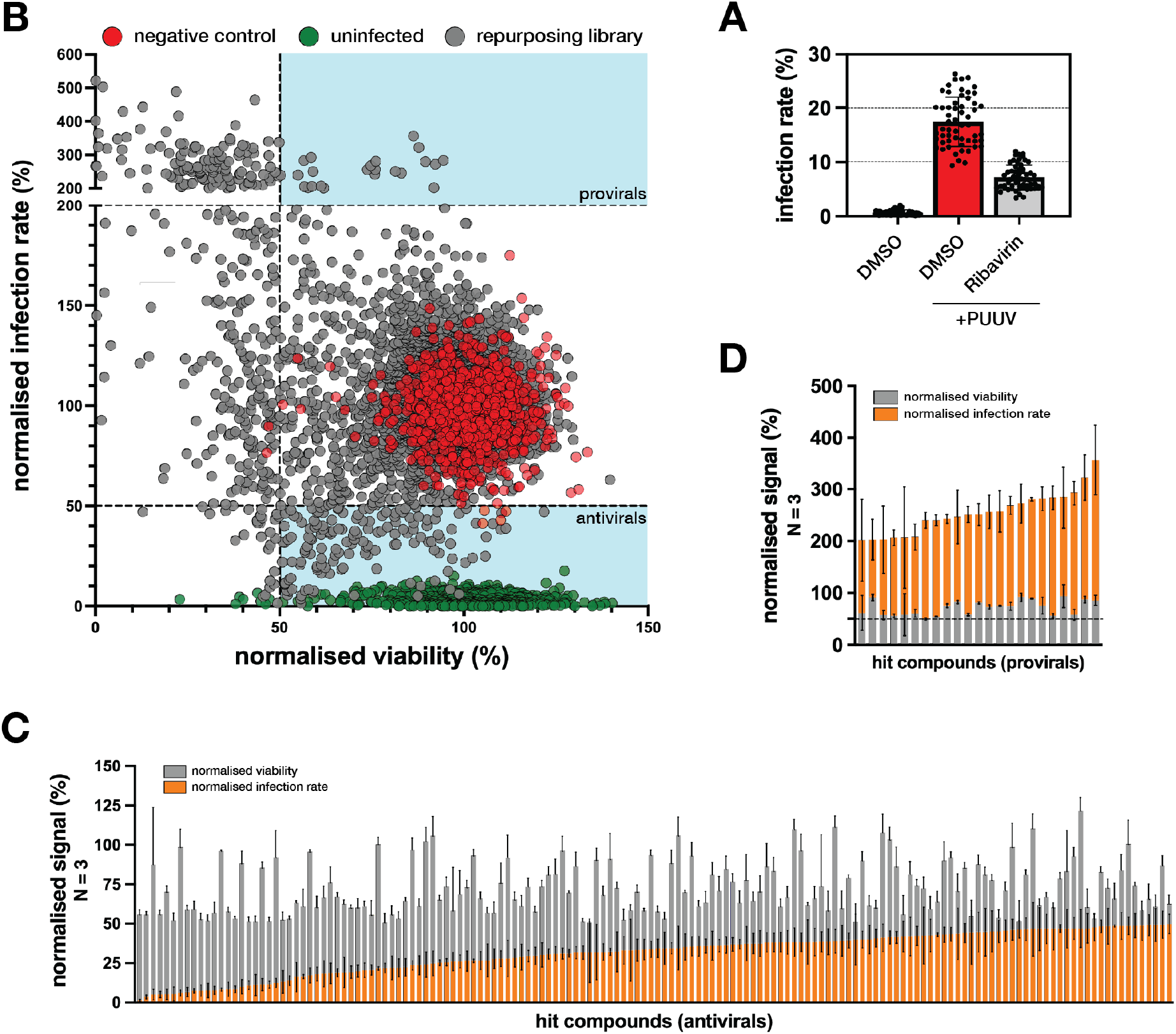
Primary screening data. A. Bar graph showing raw infection rate in the control samples. Each data point represents 1 screening plate (14 samples per plate); bars indicate mean ± SD. B. Scatter plot of the primary screening results showing normalised infection rate versus normalised viability. Grey dots represent screened compounds, red dots indicate vehicle control, and green dots indicate uninfected controls. Dashed line denote thresholds for hit calling. Shaded areas indicate regions classified as hits. C. Stacked bar graph of normalised viability (grey) and normalised infection rate (orange) for all candidate antivirals. Bars indicate mean ± SD from triplicate screening plates. D. Stacked bar graph of normalised viability (grey) and normalised infection rate (orange) for all candidate provirals. Bars indicate mean ± SD from triplicate screening plates.

### Validation of primary screening hits in A549 cells

We next carried out a comprehensive validation of the hits from the primary screen. We tested each of the 150 antiviral and 25 proviral candidates at 3 different doses, including the screening dose (**Figure 3A**). Primary screening and validation data showed a moderate but significant correlation in A549 cells (r = 0.45), indicating reproducibility across independent experiments. A substantial subset of compounds showed similar antiviral activity across experiments, while others retained a clear but attenuated effect (**Figure 3B**). Among the proviral compounds 21 out of 23 primary hits were validated in A549 cells (**Figure 3C**).

**Figure 3.**
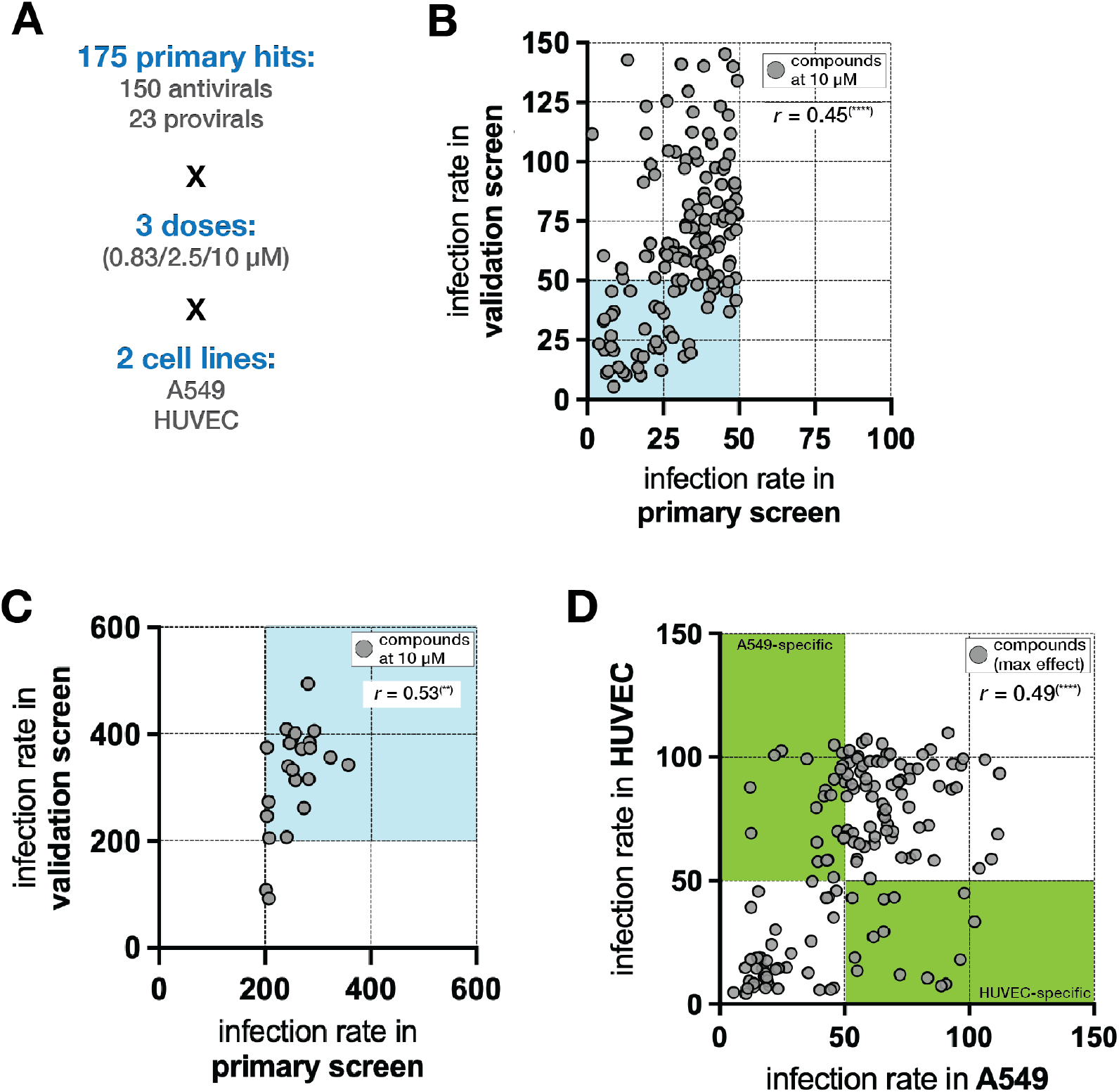
Validation of primary screening hits and cross-cell comparison. A. Overview of the validation strategy. All pro- and antiviral candidates were tested at 3 doses and in A549 cells and HUVECs. B. Scatter plot showing correlation between infection rate measured in the primary screen and in the validation experiment for antiviral compounds in A549 cells at 10 µM. Each dot represents one compound (mean of triplicates). Pearson correlation coefficient, *r*, is indicated. Shaded area indicates validated hits. C. Scatter plot showing correlation between infection rate measured in the primary screen and in the validation experiment for proviral compounds in A549 cells at 10 µM. Each dot represents one compound (mean of triplicates). Pearson correlation coefficient, *r*, is indicated. Shaded area indicates validated hits. D. Comparison of maximal antiviral effect between A549 cells and HUVECs. Each dot represents one compound; infection rates correspond to the concentration with the strongest observed effect. Pearson correlation coefficient, *r*, is indicated. Shaded areas indicate common (orange) or cell line-specific (green) hits.

### Validation of primary screening hits in HUVECs

Given that hantaviruses primarily target endothelial cells, we next tested the screening hits in a primary endothelial cell system using HUVECs (primary human umbilical vein endothelial cells), which worked robustly in our assay (**Figure S2C**). We observed substantial concordance between the datasets obtained in A549 cells and HUVECs (**Figure 3D**), with several compounds showing comparable activity levels in both systems. Nevertheless, several compounds exhibited clear cell line-specific effects. In total, 70 of the 150 primary screening hits were validated: 24 were specific to A549 cells, 10 to HUVECs, and 36 were active in both cell lines.

Due to a high infection rate in HUVECs, we could not thoroughly investigate for proviral effects in this system (**Figure S2D**).

### Functional clustering of hit compounds

One advantage of drug repurposing is the availability of prior information about the compounds, including their annotated targets and mechanisms of action. We extracted official annotations and grouped compounds into higher-level classes; missing annotations were completed by literature curation. This analysis revealed clustering of validated hits into several compound classes (**Figure 4A**), including mTOR inhibitors, HSP90 chaperone inhibitors, and compounds affecting cell signalling or nucleotide synthesis. Cross-referencing functional classes with cell type-specificity showed that most compound classes were active in both A549 cells and HUVECs. An exception was mTOR inhibitors, which were particularly effective at inhibiting infection in A549 cells (**Figure 4B**). Notably, several non-ribosome-targeting antibiotics were also present among the validated compounds. To illustrate the range of dose-response behaviours observed within functional classes, we show dose-response profiles for selected example compounds across the identified clusters (**Figure 4C**). Dose-response curves for all validated antiviral compounds are shown in **Figure S3**.

**Figure 4.**
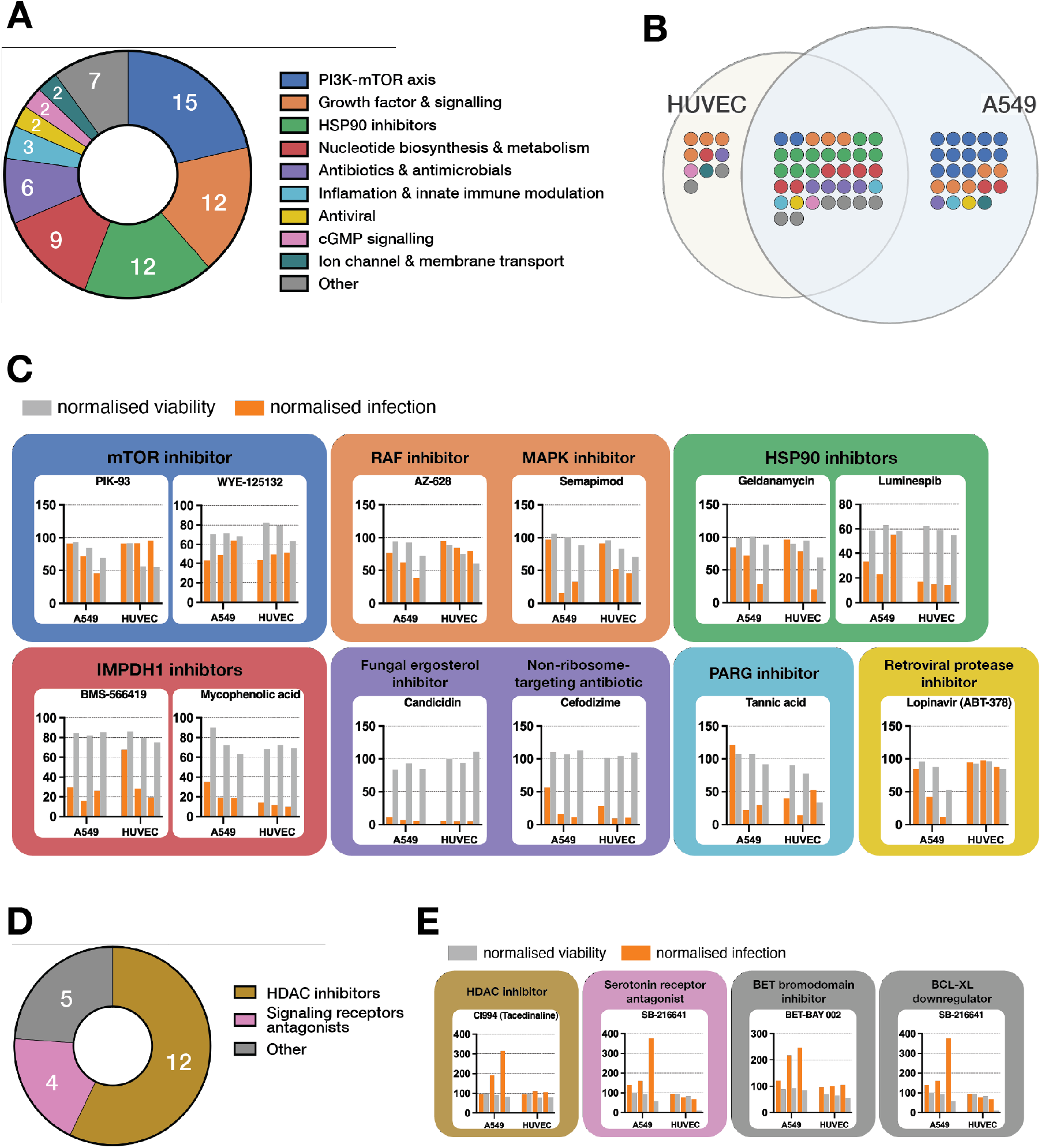
Functional classification of validated hit compounds. A. Functional distribution of validated antiviral compounds grouped according to annotated targets or mechanisms of action. Compounds were binned into higher-level categories based on Drug Repurposing Hub annotations and literature curation. Numbers indicate the number of compounds per class. B. Venn diagram showing overlap of antiviral compounds between A549 cells and HUVECs. Each dot represents one compound; colours correspond to functional classes shown in panel A. C. Representative dose-response (0.83, 2.5, 10 µM) profiles for selected antiviral compounds from major functional classes. Normalised infection rate (orange) and normalised viability (grey) are shown for A549 cells and HUVECs. Data for all validated antiviral compounds are provided in Supplementary Figure S3. D. As in A, but for candidate provirals. E. As in C, but for candidate provirals. Data for all validated proviral compounds are provided in Supplementary Figure S4.

Finally, functional clustering of the compounds that increased the infection rate in A549 cells revealed enrichment of epigenetics modifying compounds, particularly HDAC inhibitors (**Figure 4D**). The response profile of selected compounds in this group are shown in **Figure 4E**. Dose-response curves for all validated proviral compounds are shown in **Figure S4**.

Taken together, the reproducibility of antiviral effects across experiments in A549 cells and HUVECs, combined with non-random clustering of hits into functional classes, supports the internal consistency of the screening results and pin-points possible targets and signalling pathways for future repurposed drug therapeutic strategies.

## Discussion

Hantaviruses, including Puumala virus (PUUV), cause severe human diseases, yet no approved antiviral therapies are currently available and patient management remains largely supportive[3] . Although multiple targeted antiviral strategies have been explored, systematic investigations of host dependencies during productive hantavirus infection has remained limited. Here, we addressed this gap by applying a phenotypic, live-virus screening approach to identify modulators of PUUV-infection.

The output of the screen is broadly concordant with existing literature. A substantial fraction of validated antiviral compounds fall into functional classes that have been implicated previously in the replication of diverse RNA viruses, supporting the biological relevance of the screening platform. At the same time, the scale of the dataset and its validation across two distinct human cell types, A549 and HUVECs, provide insights into host dependencies difficult to achieve in hypothesis-driven studies.

Among the most robust findings is the enrichment of compounds targeting nucleotide biosynthesis, particularly inhibitors of inosine monophosphate dehydrogenase (IMPDH). IMPDH inhibitors such as mycophenolic acid and its derivatives have been reported to exhibit broad antiviral activity *in vitro*, including against hantaviruses in independent screens[12,19] . Our identification of multiple IMPDH inhibitors that suppress PUUV-infection in both A549 cells and HUVECs is therefore consistent with prior work and show that nucleotide availability is important for hantavirus replication.

Inhibitors of mTOR signalling constitute another recurrent class of antiviral compounds identified in this study. mTOR has been implicated in the life cycle of numerous RNA viruses (reviewed in[20] ). The preferential efficacy of mTOR inhibitors in A549 cells compared with HUVECs adds an additional layer of context dependence. This observation suggests that the impact of mTOR modulation on PUUV infection may depend on baseline cellular state, or on how different cell types couple translational control to antiviral responses.

Several Heat shock protein (HSP) inhibitors, particularly those targeting HSP90, also emerged among validated hits. The involvement of HSPs in viral replication is well documented across virus families, where they can facilitate protein folding, complex assembly, and stress adaptation[21] . The enrichment of HSP inhibitors in our dataset supports a role for proteostasis networks in hantavirus infection.

Unexpectedly, among the most effective and least toxic hits, we identified several beta-lactam antibiotics. While such compounds are not classically associated with antiviral effects, various classes of antibiotics have been shown to impair infection of diverse virus families, mostly through host-directed mechanisms[22] . To our knowledge, antiviral activity of beta-lactam antibiotics has not been reported previously. Given the reproducibility of our findings and the identification of multiple compounds from this class, this observation warrants further investigation.

Apart from various antiviral compounds, we identified 21 compounds with proviral effects in A549 cells, among which epigenetic modifiers, particularly histone deacetylases inhibitors, constituted the most enriched cluster. HDAC inhibitors have been reported to have both pro- and antiviral effects, depending on the virus[23,24] . Hantaviruses establish persistent infection in natural hosts[25], and viral RNA has been detected many years after symptomatic disease in a previous HPS-patient[26], suggesting hantavirus can establish long-term infection also in humans. Prior studies have linked chromatin state to viral latency and reactivation in other systems[27], future studies addressing the mechanism behind HDAC-inhibitor mediated proviral effects on PUUV infection is needed to establish a possible role for HDAC in restricting hantavirus replication or infection.

The proviral effect could only be verified in A549 cells as the higher infection rate in HUVECs compared to A549 cells (∼80% vs. ∼20%), made it difficult to analyze for pro-viral effects in HUVECs. Further studies are therefore needed to establish if the candidate provirals have cell type-specific or general effects.

Taken together, our drug repurposing data are largely concordant with existing knowledge on host-directed antiviral strategies in the hantavirus field. At the same time, our work highlights novel host dependencies during hantavirus infection that remain largely unexplored. As such, it provides a reference framework for prioritising host pathways for further mechanistic studies and for exploring new therapeutic avenues.

## Materials and Methods

### Cell lines

Human lung epithelial A549 cells (ATCC CLL-185) were grown in minimal essential medium (MEM) supplemented with 7.5% fetal bovine serum (FBS), 100 U/ml penicillin, and 100 μg/ml streptomycin. Primary human umbilical vein endothelial cells (HUVECs) (Lonza, C2517A) were grown in endothelial cell medium (ScienCell, 1001) supplemented with 5% FBS, 1% endothelial cell growth supplement (ScienCell, 1052), and 1% penicillin/streptomycin solution. Cells were incubated at 37°C and with 5% CO_2_.

### Puumala virus

PUUV (strain CG1820) was propagated on Vero E6 cells and titrated as described earlier[28] .

### Screening

For the primary screen, A549 cells were seeded onto 384-well plates (Falcon 353962) that had been pre-spotted with the Drug Repurposing Hub library. Final concentration of library compounds was 10 µM (in triplicates); DMSO was used as a negative control.

For the validation screen, A549 cells or HUVECs were seeded onto 384-well-plates pre-spotted with 150 compounds from the Drug Repurposing Hub library. Final concentrations of these compounds were 0.83 µM, 2.5 µM and 10 µM (in triplicates); DMSO was used as a negative control. After 6 hours incubation of cells and compounds, PUUV was added to the wells. After a further 24-hour incubation, cells were fixed and directly stained for microscopy, as described below.

### Immunofluorescence microscopy

Cells were fixed with 4% formaldehyde for 15 minutes at room temperature, followed by 10-min permeabilization with 0.1% Triton X-100/PBS, 30 min blocking with blocking buffer (3% BSA/0.1% Tween-20/PBS), overnight incubation with primary antibody (convalescent patient sera; 1:1000) at 4°C, and a 1-h incubation with secondary antibody (Invitrogen, A-21445) solution supplemented with 2 µM Hoechst 3342 (Thermo Scientific, 62249) and 1 µM CellTracker Orange CMTMR Dye (Thermo Scientific, C2927). PBS washes were done between each step.

Cells were then imaged using an IN Cell Analyzer 2200 (GE Healthcare) high-throughput microscope. Image analysis was performed using CellProfiler[18] and data processing was done using the KNIME Analytics Platform.

## Acknowledgments

The authors want to thank the Chemical Biology Consortium Sweden for providing the compound library. Work done in O.F.-C. lab was supported by grants from Swedish Research Council (E0003101, 2023-02532). Work done in J.K. lab was supported by grants from Swedish Research Council (2024-02578).

## Author contribution

W.C. and B.P. conceived the study, performed all experiments, and wrote the manuscript; B.P. analysed the data and prepared the figures; O.F.-C. and J.K. provided funding and supervised the study. All authors read and revised the manuscript.

## Data availability statement

Primary screening data and library metadata are available in the Supplementary information.

## Figure legends

**Figure S1.**
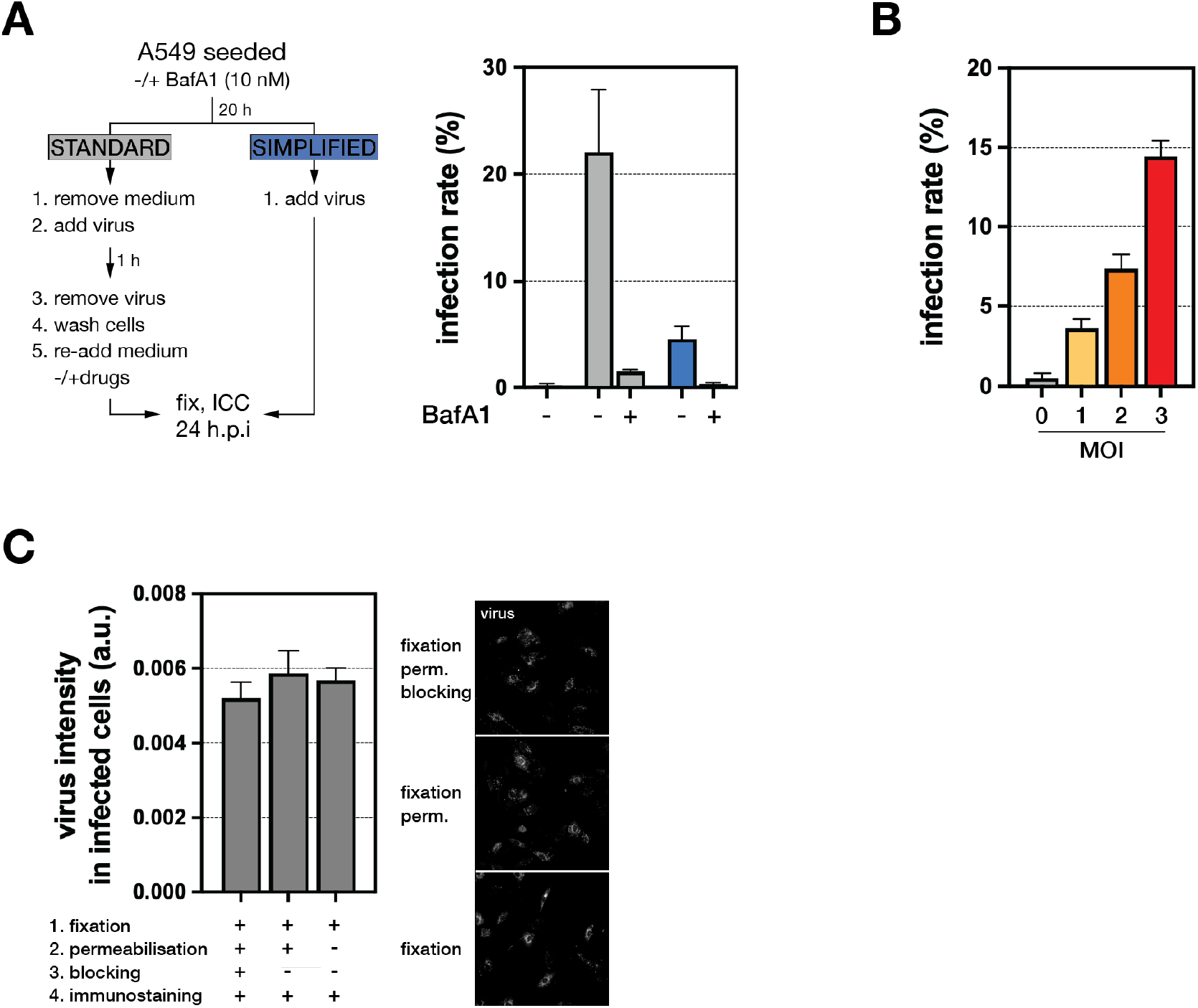
Simplification and optimisation of the screening assay. A. Comparison of standard and simplified infection protocols. In the standard protocol, growth medium was removed, virus was added for 1 h, then removed, cells were washed, and fresh medium with or without compounds was added. In the simplified protocol, virus was added directly to the growth medium, without removal and washing steps. Bar graph shows quantification of the infection rate. Bars indicate mean ± SD (N = 3); colours correspond to the infection protocol. B. Optimisation of multiplicity of infection (MOI). A549 cells were infected with increasing MOI (0-3). Bar graph shows the raw infection rate. Each bar indicates mean ± SD (N = 3). C. Simplification of the immunostaining protocol. Bar graph (left) shows quantification of the virus signal in infected cells following different immunostaining protocols. Representative images are shown on the right. Each bar indicates mean ± SD (N = 3).

**Figure S2.**
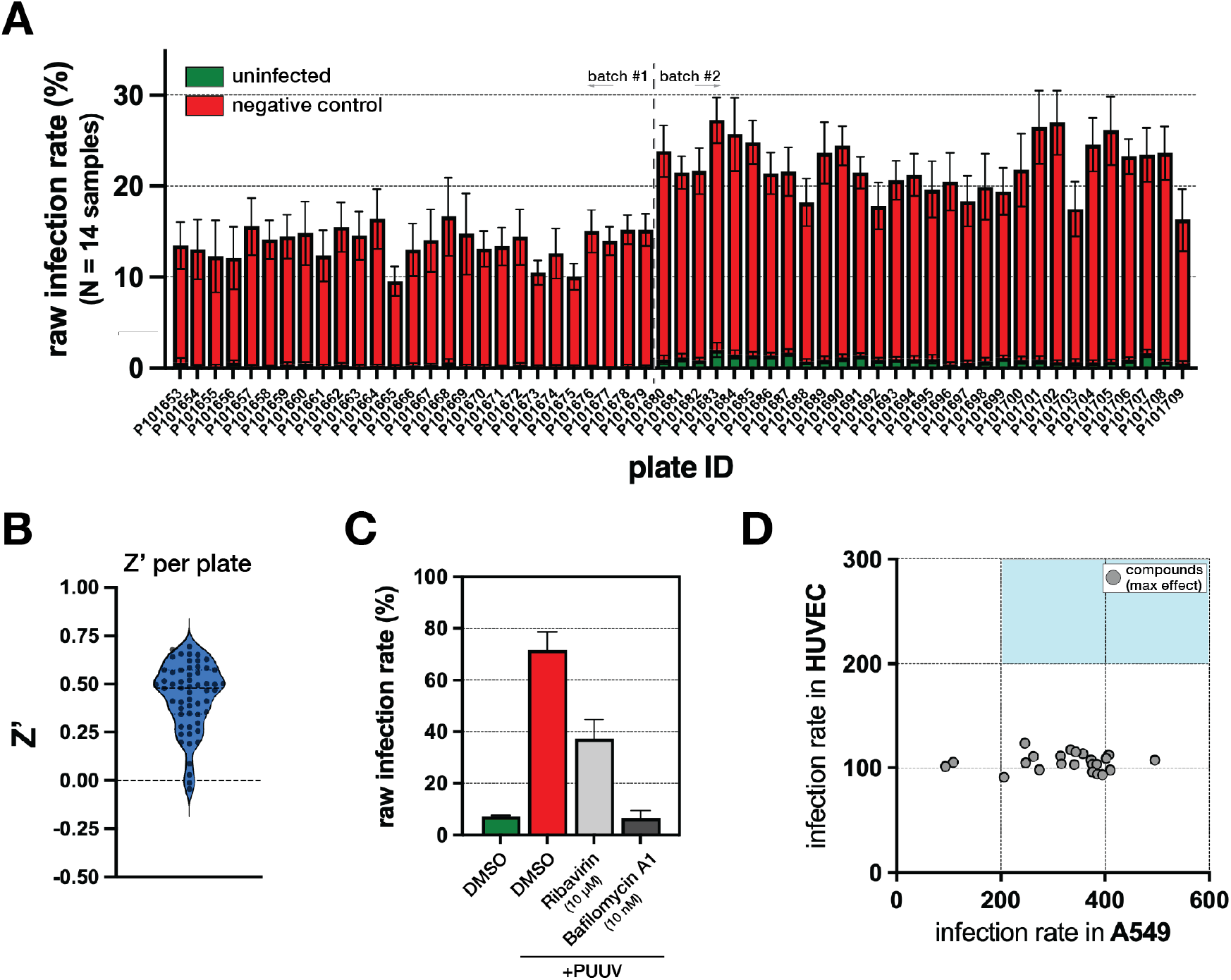
Primary screening quality control metrics. A. Bar graph showing raw infection rate in uninfected (green) and infected vehicle-treated (red) samples. Each bar indicates mean ± SD (n = 14) in one screening plate. Dashed vertical line demarcates screening batches. B. Violin plot showing the distribution of Z’ values calculated for each plate (dot). Mean Z’ for the entire screen was 0.41. C. Quantification of PUUV infection in HUVECs following treatment with vehicle (DMSO), 10 µM Ribavirin, or 10 nM Bafilomycin A1. Infection rate was calculated as the percentage of virus-positive cells. Dots represent individual wells, each with 2000-2500 cells. Bars indicate mean ± SD. D. Comparison of maximal proviral effect between A549 cells and HUVECs. Each dot represents one compound; infection rates correspond to the concentration with the strongest observed effect. Pearson correlation coefficient, *r*, is indicated. Shaded areas indicate common (orange) or cell line-specific (green) hits.

**Figure S3.**
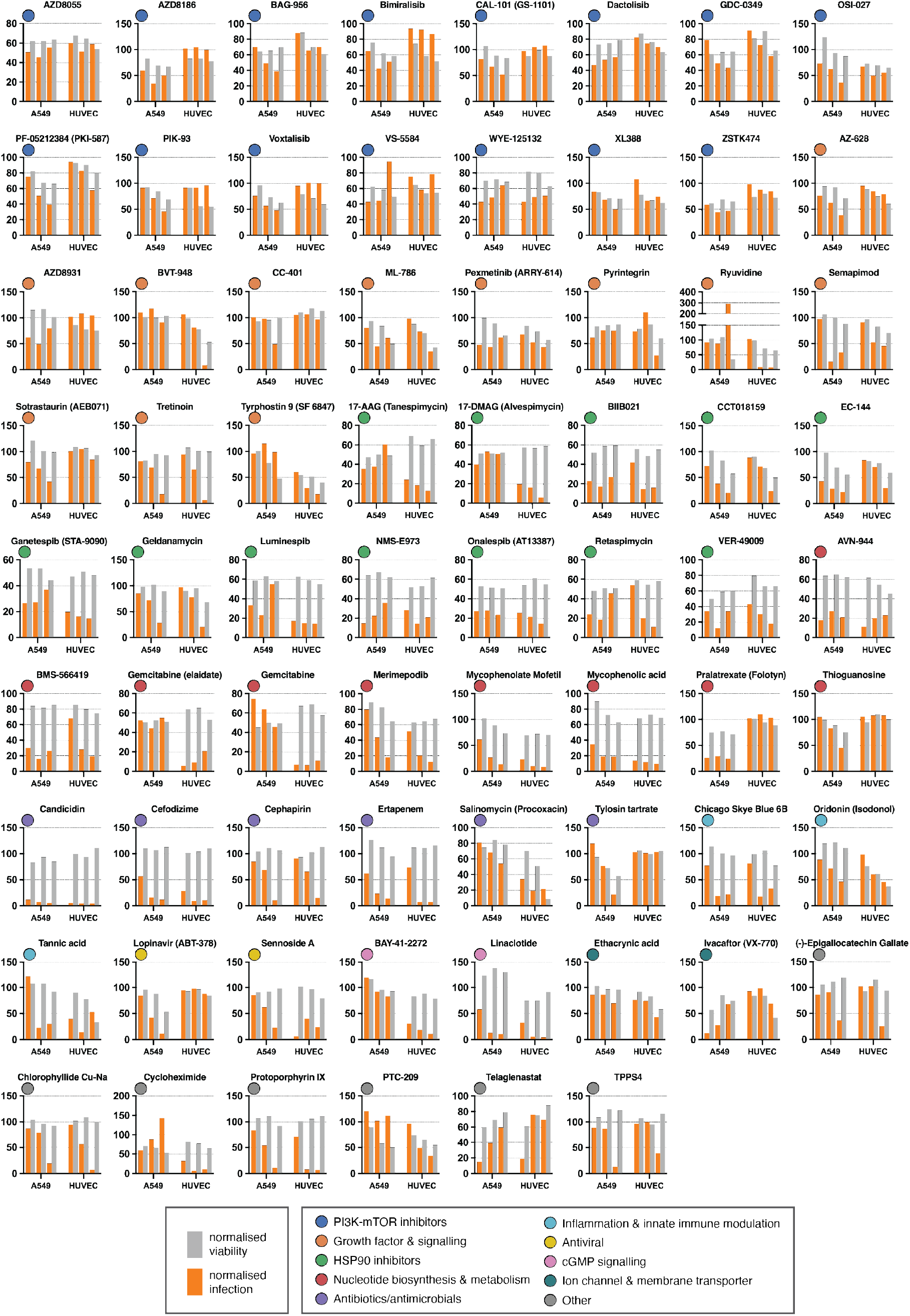
Dose response curves of validated antivirals. A. Dose response profiles for all validated antiviral compounds. Normalised infection rate (orange) and normalised viability (grey) are shown for A549 cells and HUVECs. Coloured circles correspond to functional classes.

**Figure S4.**
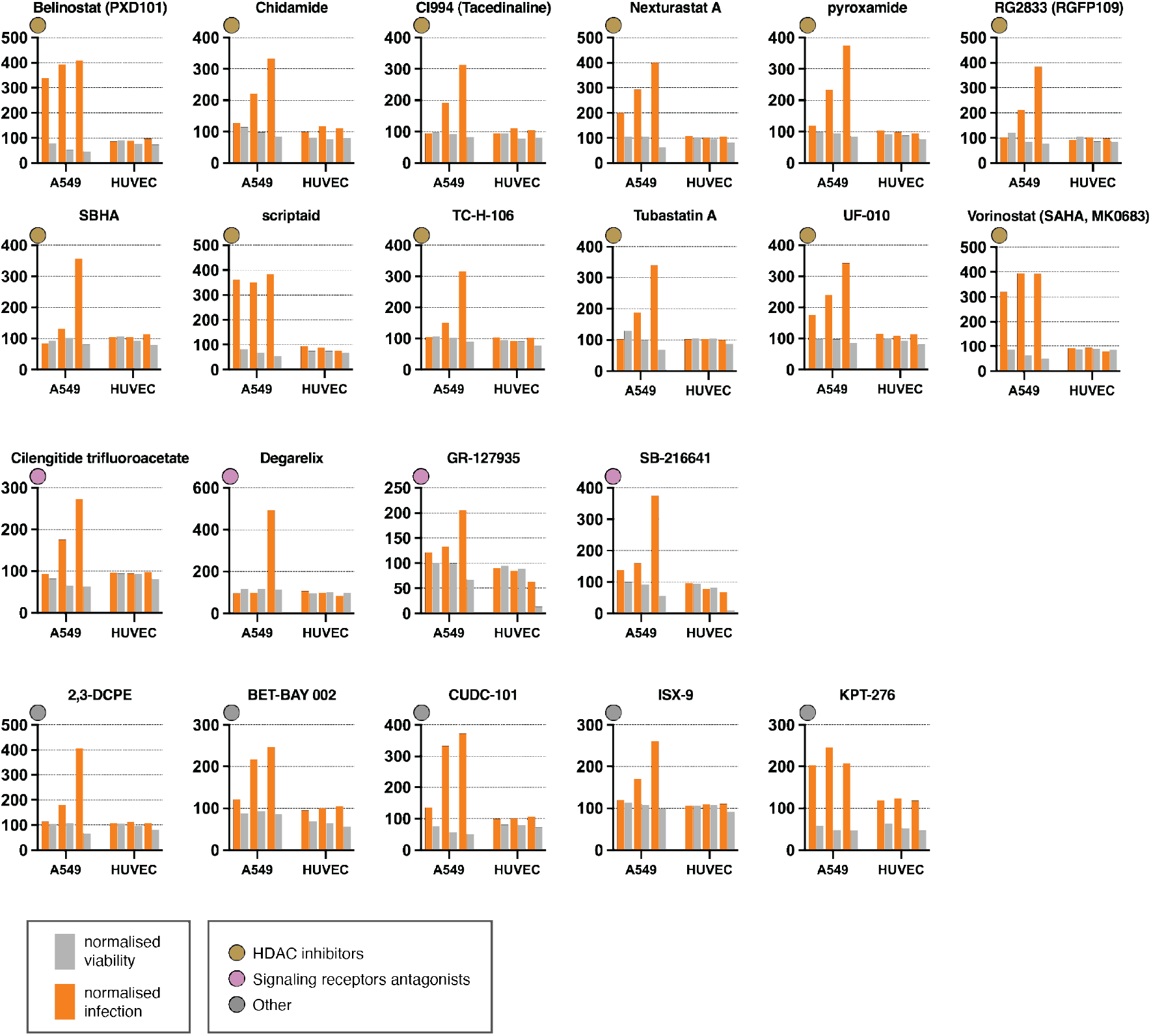
Dose response curves of validated provirals. A. Dose response profiles for all validated proviral compounds. Normalised infection rate (orange) and normalised viability (grey) are shown for A549 cells and HUVECs. Coloured circles correspond to functional classes.

